# Preservation of cerebral vascular and metabolic health with greater cardiorespiratory fitness in coronary artery disease

**DOI:** 10.1101/2025.08.22.671781

**Authors:** Safa Sanami, Stefanie A. Tremblay, Zacharie Potvin-Jutras, Ali Rezaei, Dalia Sabra, Christine Gagnon, Brittany Intzandt, Amélie Mainville-Berthiaume, Lindsay Wright, Mathieu Gayda, Josep Iglesies-Grau, Anil Nigam, Louis Bherer, Claudine J. Gauthier

## Abstract

**Background:** Coronary artery disease (CAD) is the leading cause of cardiovascular-related death globally. Beyond its cardiac consequences, CAD significantly impacts brain health, causing grey matter atrophy, reduced cerebral blood flow (CBF), impaired cerebrovascular reactivity (CVR), and cognitive decline. Recent evidence also highlights altered cerebral metabolism in CAD, characterized by reduced cerebral metabolic rate of oxygen consumption (CMRO₂) and increased oxygen extraction fraction (OEF). Notably, impaired cardiorespiratory fitness, as measured by reduced peak oxygen uptake (VO₂peak), is a hallmark of CAD progression and a strong prognostic marker of cardiovascular and neurological outcomes. Although VO₂peak is associated with brain structural integrity and cognitive function, its relationship to cerebral vascular and metabolic function in CAD patients remains poorly understood.

**Methods:** Thirty-seven healthy individuals (age=65.35 ± 8.31) and thirty-five patients with CAD (age=66.42 ± 9.29) participated in this study. The objective was to determine whether higher cardiorespiratory fitness is associated with markers of cerebral health known to be impaired in CAD, in order to assess the potential of exercise as a strategy to mitigate CAD-related brain alterations. Cerebral vascular and metabolic biomarkers, including CBF, CVR, CMRO₂, and OEF, were quantified using calibrated functional magnetic resonance imaging (MRI). Participants also completed a maximal cardiopulmonary exercise test on a bicycle ergometer to determine peak oxygen uptake (VO₂peak).

**Results:** Across all participants, VO₂peak was positively associated with CBF (β=0.32, p=0.02), and CVR (β=0.002, p=0.04), in grey matter, confirming a link between aerobic fitness and vascular health across the cardiovascular health spectrum. However, metabolic markers exhibited group-specific patterns. Specifically, in CAD patients, VO₂peak was positively associated with CMRO₂ (*β* = 0.08, *p* = 0.02), suggesting that reduced CMRO₂ in CAD may be partially preserved by greater cardiorespiratory fitness. In contrast, a negative association between VO₂peak and OEF was observed exclusively in healthy controls (*β* = −3.6, *p* = 0.02), consistent with adaptations in healthy aging being primarily driven by improved CBF without changes in CMRO₂.

**Conclusion:** This study shows that higher cardiorespiratory fitness is associated with improved cerebral vascular and metabolic function, with distinct patterns observed between healthy individuals and those with CAD. In patients with CAD, greater fitness appears to preserve cerebral oxygen metabolism, while in healthy individuals, fitness is primarily linked to enhanced perfusion. These findings support the role of aerobic exercise as a promising strategy to counteract CAD-related brain alterations, emphasizing the importance of targeting cardiorespiratory fitness in both prevention and rehabilitation settings.

## Introduction

Coronary artery disease (CAD), caused by atherosclerotic narrowing of coronary arteries is the most common form of cardiovascular disease and first cause of death worldwide (Stark et al., 2024). In addition to increased mortality risk, CAD also has extensive effects beyond cardiac health, significantly impacting the brain. CAD is associated with grey matter atrophy, reduced cerebral blood flow (CBF), impaired cerebrovascular reactivity (CVR), which is the ability of brain vessels to dilate or constrict appropriately in response to metabolic demands (Almeida et al., 2008; Anazodo et al., 2016). Collectively, these vascular and structural impairments contribute to an elevated risk of cognitive decline (Rostamian et al., 2015; Xia et al., 2020). In our recent work (Sanami et al., 2025), we demonstrated that individuals with CAD exhibit not only deficits in CBF and CVR, but also altered brain metabolism, with individuals with CAD showing reduced cerebral metabolic rate of oxygen consumption (CMRO₂)—the rate at which brain tissue consumes oxygen to sustain neuronal activity—and elevated oxygen extraction fraction (OEF), a marker of the balance between oxygen delivery and consumption, reflecting the proportion of oxygen extracted from blood by brain tissue.

Structured cardiac rehabilitation programs aimed at improving cardiorespiratory fitness improve patient survival and quality of life in cardiovascular disease, with VO₂peak levels and their improvement being strong independent predictors of morbidity and mortality in CAD (Bocalini et al., 2008; Carbone et al., 2022). Beyond cardiovascular improvements, higher cardiorespiratory fitness—typically measured by peak oxygen uptake (VO₂peak)—is associated with beneficial effects on brain structure and function. For instance, improved VO₂peak following rehabilitation correlates positively with increased brain volume (MacIntosh et al., 2014). Additionally, cardiac rehabilitation has been shown to directly enhance CBF, suggesting beneficial cerebrovascular adaptations in CAD (Anazodo et al., 2016; MacIntosh et al., 2014). However, the relationship between VO₂peak and other affected brain physiological components in CAD, such as CVR and metabolic function, remains almost completely unexplored.

In normal aging, VO₂peak has been positively associated with better executive function and memory (Erickson et al., 2011; Hötting and Röder, 2013), as well as with greater brain volume in the frontal and temporal lobes and the hippocampus (Erickson et al., 2009; Weinstein et al., 2013). However, its relationship with physiological markers—such as CBF, CVR, CMRO₂, and OEF—is less well documented. The relationship between VO₂peak and vascular biomarkers such as CBF and CVR has been documented in a few populations. CBF was found to be positively associated with VO₂peak in children (Chaddock-Heyman et al., 2016), young and older adults (Intzandt et al., 2020), CAD patients (MacIntosh et al., 2014) and in mild cognitive impairment (MCI) (Maffei et al., 2017). These studies typically show positive associations between VO₂peak and CBF, suggesting that greater aerobic fitness may enhance cerebral perfusion and support brain function (Chaddock-Heyman et al., 2016; MacIntosh et al., 2014; Maffei et al., 2017). However, a few studies have documented null or even negative associations—particularly in older populations—suggesting that age may moderate the relationship (Intzandt et al., 2020; Olivo et al., 2021).

Only a few studies have investigated the relationship between VO₂peak and CVR, showing a more complex relationship with fitness than CBF. (DuBose et al., 2022) have suggested that the relationship may follow an inverse U-shape, with a positive relationship between CVR and VO₂peak for individuals who are less physically active and a negative relationship for those who are more active than average. This is consistent with several studies in the aging literature(Intzandt et al., 2020; Potvin-Jutras et al., 2024; Thomas et al., 2014), but the relationship in populations with heart disease remains to be established. Given that both CBF and CVR are affected in CAD (Anazodo et al., 2016) and show a relationship with cardiorespiratory fitness in normal aging, examining their relationship with VO₂peak in CAD may provide insight into mechanisms linking cardiorespiratory fitness to brain vascular health in this population.

While the relationship between cardiorespiratory fitness and CBF has often been reported, there is a lack of research on how cardiorespiratory fitness affects resting cerebral metabolic biomarkers (CMRO_2_ and OEF). This gap persists despite evidence from functional near-infrared spectroscopy (fNIRS) studies suggesting that higher fitness levels may influence brain oxygenation dynamics during exercise and cognitive tasks, particularly in older adults (Agbangla et al., 2019; Dupuy et al., 2015). To date, only two recent studies have directly explored the relationship between VO₂peak and brain metabolism, both focusing exclusively on healthy older adults. (Huppert and Qiao, 2025) reported a negative relationship between global CMRO₂ and VO₂peak, whereas a recent study from our group (Sanami & Rezaei et al., 2025) found no significant association between VO₂peak and CMRO₂. Instead, this previous study identified a negative relationship between VO₂peak and oxygen extraction fraction (OEF), highlighting potential differences in how fitness impacts cerebral metabolism and oxygen availability. However, this relationship has not been explored in CAD, where metabolic impairments may be more pronounced (Sanami et al., 2025). This study aims to comprehensively evaluate—for the first time—the association between VO₂peak and cerebral metabolic markers (CMRO₂ and OEF) across the whole brain in both CAD patients and healthy controls. Documenting these relationships could help identify sensitive metabolic biomarkers linked to fitness, facilitating more targeted interventions to preserve brain health in clinical populations.

Therefore, the aim of this study is to investigate the relationships between VO₂peak and a comprehensive array of cerebral vascular and metabolic biomarkers in individuals with CAD as compared to healthy controls (matched by age and sex). We hypothesize that greater VO₂peak will be associated with better cerebral vascular and metabolic health, denoted by higher CBF, CVR and CMRO_2_ and lower OEF, especially in CAD patients given their larger deficits. This study will break new ground in our understanding of how cardiorespiratory fitness can improve brain physiological health in CAD patients.

## Materials and methods

### Participants

Ninety-nine (99) participants (16 females) of 50 years old and above were recruited, from which 87 completed the study (46 CAD patients and 41 healthy controls; HC). Out of 12 participants that discontinued their participation, five participants did not complete the study due to interruptions during the COVID-19 pandemic, two participants were excluded due to claustrophobia, two participants were excluded due to discomfort during the MRI and three participants chose not to participate for personal reasons (e.g., loss of interest, issues with scheduling). Of the 87 remaining participants, twelve had only 5 minutes of resting CBF data instead of the full acquisition including hypercapnia, due to discomfort during hypercapnia, and these were included in CBF analysis only. Seventy-five (75) participants had a complete dataset (Figure 1 for the flowchart). To ensure sufficient data quality, we excluded three participants with low-quality imaging data. Details for exclusion will be given in the preprocessing section. The final sample included 35 CAD patients and 37 control participants with gas challenge (Table 1 for demographics). The study received approval from the Comité d’éthique de la recherche et du développement des nouvelles technologies (CÉRDNT) of the Montreal Heart Institute in accordance with the Declaration of Helsinki. Data collection was conducted at the Montreal Heart Institute.

**Figure 1.**
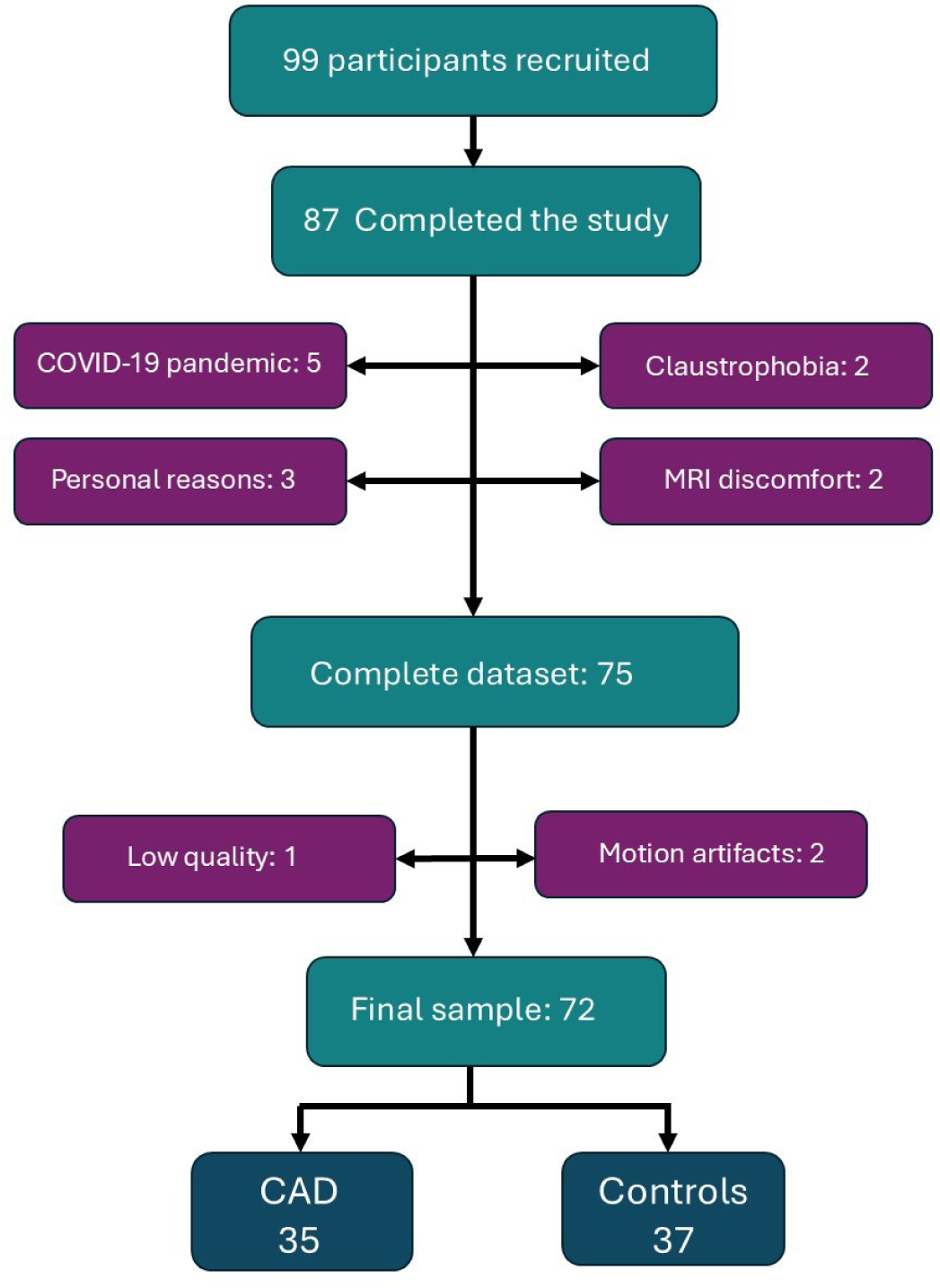
Participant Recruitment and Final Sample Selection. Flowchart showing participant recruitment and retention. A total of 99 participants aged ≥50 years were recruited. Twelve discontinued participation due to various reasons, including the COVID-19 pandemic, claustrophobia, MRI discomfort, and personal reasons. Of the 87 who completed the study, 75 had complete datasets. After excluding three participants for low-quality data or excessive motion, the final sample included 72 individuals (35 CAD patients and 37 controls).

**Table. 1.** Demographic data on participants with gas challenge data.

|  | **Controls (37)** | **CAD (35)** | **P-value** |
| --- | --- | --- | --- |
| Age (years) | 65.35 ± 8.31 | 66.42 ± 9.29 | 0.83 |
| Sex | Female (10) | Female (6) | - |
| Education (years) | 16.22 ± 2.45 | 16.03 ± 4.03 | 0.80 |
| VO<sub>2</sub>peak<sup>a</sup> (ml/min/kg LBM) | 27.95 ± 6.46 | 22.14 ± 5.11 | **<0.01** |
| MoCA<sup>b</sup> | 26.73 ± 2.01 | 25.33 ± 3.04 | **0.02** |
| MMSE<sup>c</sup> | 28.25 ± 1.48 | 28.42 ± 1.17 | 0.97 |
| Executive Function(z-score) | 0.314 ± 0.62 | -0.32 ± 1.01 | **<0.01** |
| Working Memory (z-score) | 0.04 ± 0.83 | 0.033 ± 0.79 | 0.95 |
| Processing Speed (z-score) | 0.26 ± 0.72 | -0.34 ± 0.86 | **<0.01** |
| Verbal/Episodic Memory (z-score) | 0.19 ± 0.96 | 0.041 ± 0.97 | 0.31 |
| SBP (mmHg)<sup>d</sup> | 127.12 ± 13.85 | 126.96 ± 14.54 | 0.96 |
| BMI (kg/m<sup>2</sup>)<sup>e</sup> | 26.11 ± 3.94 | 26.95 ± 3.57 | 0.36 |

<sup>a</sup>VO<sub>2</sub>peak: Peak oxygen uptake during maximal exercise; a measure of cardiovascular fitness.
<sup>b</sup>MoCA : Montreal Cognitive Assessment; a 30-point test screening for mild cognitive impairment.
<sup>c</sup>MMSE: Mini-Mental State Examination; a brief 30-point test to assess global cognitive function.
<sup>d</sup>SBP: Systolic Blood Pressure
<sup>e</sup> BMI: Body Mass index

Inclusion criteria for CAD patients required a confirmed history of coronary artery disease, including prior acute coronary syndromes—such as unstable angina, non–ST-segment elevation myocardial infarction (NSTEMI), or ST-segment elevation myocardial infarction (STEMI)—as well as prior coronary revascularization with either percutaneous coronary intervention (PCI) or coronary artery bypass grafting (CABG), or clinically documented angina. Coronary angiography was performed in most cases as part of standard diagnostic care to guide clinical management (Table 2 for clinical characteristics). Healthy controls (HCs) were free of cardiac disease, diabetes, hypertension and cognitive impairment. All participants had to be fluent in English or French to ensure informed consent and accurate cognitive assessment.

**Table. 2.** Clinical information in CAD and HC participants Clinical Health characteristics in patients with CAD (n=35), Healthy Controls (n=37)

| **Clinical Health characteristics in patients with CAD (n=35), Healthy Controls (n=37)** |  |  |
| --- | --- | --- |
| **Cardiometabolic risk factors , n(%)** |  |  |
|  | Patients with CAD | Healthy Controls |
| Diabetes | 3 (8) | - |
| Hypertension | 18 (51) | - |
| Dyslipidemia | 28 (80) | 3(8) |
| Obesity | 5(14) | 5(11) |
| Familial history of cardiovascular disease | 21 (60) | 6(16) |
| **Cardiovascular history, n(%)** |  |  |
| Previous MI<sup>a</sup> | 14 (40) | - |
| STEMI<sup>b</sup> | 8 (22) | - |
| NSTEMI<sup>c</sup> | 6 (17) | - |
| Percutaneous coronary intervention (PCI) | 22 (62) | - |
| Coronary artery bypass grafting (CABG) | 10 (28) | - |
| Angina | 10 (28) | - |
| **Cardiovascular medication, n (%)** |  |  |
| Beta-Blocker | 17 (48) | - |
| Aspirin | 11 (31) | - |
| Antilipemic agents (e.g., statins) | 26 (74) | 1(2) |
| RAAS<sup>d</sup> inhibitors | 11 (31) | - |
| Calcium channels blockers | 5 (14) | - |

<sup>a</sup>MI = Myocardial Infarction
<sup>b</sup>STEMI = ST-segment elevation Myocardial Infarction
<sup>c</sup>NSTEMI = Non-ST-segment elevation
<sup>d</sup>RAAS = Renin-Angiotensin-Aldosterone System

Exclusion criteria for all participants included a history of stroke, neurological, psychiatric or respiratory disorder, thyroid disease, potential cognitive impairment (MMSE < 25), tobacco use, high alcohol consumption (more than two drinks per day), and contraindications to MRI (e.g., ferromagnetic implants, claustrophobia). Participants were also excluded if they had undergone surgery under general anesthesia within the past 6 months, had a recent acute coronary event (< 3 months), chronic systolic heart failure, resting left ventricular ejection fraction < 40%, symptomatic aortic stenosis, or severe non-revascularizable coronary artery disease, such as left main coronary stenosis. Other exclusion factors included awaiting coronary artery bypass surgery, having an implanted automatic defibrillator or permanent pacemaker, malignant arrhythmias during exercise, arthritis or claudication, severe exercise intolerance, or excessive discomfort due to hypercapnia (> 5 on the (Banzett et al., 1996), dyspnea scale).

Data was collected over four visits. The first visit established eligibility through a medical history questionnaire and assessed global cognitive functioning via the Mini Mental State Examination (MMSE) (Folstein et al., 1975). Additionally, participants underwent a 2-minute hypercapnia test to ensure their ability to tolerate the respiratory manipulation during the subsequent MRI session (with 2 participants being excluded at this stage). The second visit involved administration of the cognitive assessment. During the third visit, participants completed a maximum effort test to determine their VO₂peak. The final visit was dedicated to the MRI acquisition.

### Maximal cardiopulmonary exercise test (CPET)

Incremental CPET to volitional exhaustion was performed on a cycle ergometer (E100, Cosmed, Italy) according to the latest recommendations and as previously published (Guazzi et al., 2012; Kirsch et al., 2024). A 3-minute warm-up phase at an initial workload of 20 W was followed by an incremental exercise test, with increases of 10 to 20 W per minute depending on the participant’s physical capacity, performed at a pedaling cadence between 60 and 80 rpm. The recovery phase consisted of 2 minutes of active recovery at 20 W at pedalling speed between 50 and 60 rpm, followed by 3 minutes of passive recovery where participants remained seated quietly on the ergometer. Gas exchange parameters were continuously measured at rest, during exercise, and during recovery using a metabolic system (Cosmed Quark, Cosmed, Italy), capturing on a breath-by-breath basis and then averaged in 10 second increments as recently published (Kirsch et al., 2024). There was continuous Electrocardiogram (ECG) monitoring before, during the test and in the recovery (T12x, Cosmed, Italy). Diastolic (DBP) and systolic (SBP) blood pressures (Tango M2, Suntech, USA) and ratings of perceived exertion (RPE) were measured at rest and every 3 minutes throughout the test. Participants reached maximal effort when one of the following four criteria were met: (1) a plateau of the VO_2_ (<150 mL/min in the last 30 s of effort) despite increased power; (2) a respiratory exchange ratio >1.10; (3) an inability to maintain 50 rpm; or (4) patient exhaustion due to fatigue or other clinical symptoms (ECG and/or blood pressure abnormalities requiring exercise cessation (Guazzi et al., 2012).

The highest VO₂ value reached during the exercise phase, averaged over a 10-second interval, was considered VO₂peak and expressed in mL/min/kg of lean body mass (Krachler et al., 2015). Lean body mass was estimated using a body composition analyzer (BC 418, TANITA, Japan), which also provided measurements of fat mass.

### MRI acquisition

Data were collected using a 3T Skyra MRI system with a 32-channel head coil. The acquisition protocol included structural MRI, dual-echo pseudo-continuous Arterial Spin Labeling (pCASL) for simultaneous acquisition of perfusion and Bold Oxygen Level Dependent (BOLD) signals and a blood magnetization map (M0) for perfusion quantification. pCASL data was acquired with a voxel resolution of 3.4375 x 3.4375 x 7 mm, TR/TE1/TE2/alpha: 4000/10/30ms/90°, labeling duration of 1517 ms with a post-labeling delay (PLD) of 1300 ms. The M0 acquisition had identical parameters but with a TR of 10 seconds, to ensure a fully recovered magnetization.

Because the MRI data was acquired within two sub-sessions to allow the participant to take off the mask for the rest of the data acquisition, two T1-weighted acquisitions were collected to ensure registration accuracy of the pCASL slab. One lower resolution acquisition was performed right before pCASL while the person had the mask on, and a higher resolution acquisition was performed during the second sub-session, without the mask. The low resolution T1-weighted acquisition was acquired using a magnetization prepared rapid gradient echo (MPRAGE) sequence with TR/TE/Flip angle = 15 ms/3.81 ms/25° with a 1.5 mm isotropic resolution. The high-resolution T1-weighted structural images were acquired with a MPRAGE sequence, TR/TE/Flip angle = 2300 ms/2.32 ms/8° with a 0.9 mm isotropic resolution.

### Respiratory manipulation

The RespirActTM system (RespirAct_TM_, Thornhill Research, Toronto, Canada) was used during the breathing manipulation to target specific end-tidal partial pressures of CO₂ and O₂. Three conditions were sequentially targeted for 2 minutes, preceded and followed by 2 minutes of air inhalation: a hypercapnic condition (targeting 5 mmHg above baseline for end-tidal partial pressure CO₂ (PETCO₂)), an isocapnic hyperoxic condition (targeting 150 mmHg for end-tidal O₂), and a combined hyperoxia and hypercapnia condition (targeting 5 mmHg above baseline for PETCO₂ and 150 mmHg for PETO₂). A rebreathing face mask was used to deliver gases to the participants. The system delivered gas at a flow rate of 20 L/min, while the concentrations of exhaled CO_2_ and O_2_ were monitored. On their first study visit, a familiarization session was done with a 2min hypercapnia manipulation to ensure participant comfort during the MRI. Breathing discomfort was assessed using the Banzett dyspnea scale and those with a score >5 were not invited to continue in the study (n = 2)(Banzett et al., 1996).

#### Respiratory data analysis

CO_2_ and O_2_ were sampled continuously throughout the breathing manipulation by the RespirAct, which also generates a time series of end-tidal CO_2_ and O2 partial pressures. The end-tidal partial pressure is used as a proxy for arterial gas partial pressure (Slessarev et al., 2007). To combine these values with the MRI data, a MATLAB script was employed to smooth the data, remove outliers and resize the data to match the durations of the BOLD and ASL signals.

### Imaging data processing

#### T1 processing

All structural images underwent preprocessing with the Brain Extraction Tool (BET) in FSL to remove the skull. Subsequently, FSL’s FAST was employed to segment the structural images into GM, White Matter (WM), and Cerebrospinal Fluid (CSF).

#### Preprocessing of BOLD and pCASL

Both ASL and BOLD datasets were motion-corrected using FSL’s mcflirt. A brain mask was created from the motion-corrected BOLD images using BET and applied to both echoes to remove the scalp. The first echo of the dual-echo pCASL acquisition was used to compute the perfusion-weighted time-series, from which CBF was estimated via surround subtraction. Volumes with voxel intensities exceeding 3 standard deviations from the mean were flagged, and those with >50% outlier voxels were excluded to ensure sufficient temporal signal-to-noise ratio (tSNR) (Clement et al., 2022). Participants were excluded if their pCASL data showed a tSNR below 0.5 or if they exhibited excessive motion during acquisition, defined as translational head movements exceeding 2 mm, to ensure overall data quality.

The M0 image was skull-stripped using BET and used for perfusion calibration. The second echo was used to extract the BOLD-weighted signal via surround addition, reflecting changes in blood oxygenation.

#### CBF and CVR maps

Perfusion was quantified from the preprocessed ASL time-series using FSL’s BASIL with kinetic modeling and M0 acquisition. Partial volume correction was applied via weighted averaging based on tissue contributions from GM, WM, and CSF segmentations of the T1-weighted data (Asllani et al., 2008). CVR was quantified from the BOLD time-series derived from the second echo of the pCASL data using surround addition. Due to dead space in the tubing, there is an individual-specific delay between gas sampling and the corresponding brain MRI signal, requiring adjustment of end-tidal traces. This lag was estimated via cross-correlation using a custom MATLAB script, selecting the delay where PETCO2 and the mean GM BOLD signal were maximally correlated (Sleight et al., 2021). Absolute CVR was calculated as the percent BOLD signal change per mmHg CO2 from a General Linear Model (GLM) using the ETCO2 time course as the regressor.

#### OEF & CMRO_2_ maps

OEF was estimated using the general calibration model (GCM), based on the deoxyhemoglobin dilution model of the BOLD signal (Gauthier and Hoge, 2012). Detailed equations and the full estimation process for OEF are provided in the Supplementary material. Following OEF estimation, CMRO_2_ was calculated using CBF maps and arterial oxygen content using Fick’s principle, as described in Equation 3 of the supplementary material.

#### Registration to MNI space

The pCASL images were registered to MNI space in three separate steps. First, a rigid body registration with nearest neighbor interpolation was conducted using Advanced Normalization Tools (ANTs) to align the pCASL mean images with the low-resolution (1.5 mm) T1-weighted image. Next, the low-resolution T1-weighted images were registered to the high-resolution (0.9 mm) T1 space using a multistage approach that combined rigid, affine, and nonlinear (SyN) transformations with a multi-resolution strategy. In the final step, the high-resolution T1 images were registered to MNI space using the same transformation method applied for the registration from low to high resolution. Finally, the pCASL images were registered to MNI space by applying the three transformation matrices obtained from the previous registration steps, utilizing 3 warp images.

#### Statistical Analysis

To examine whether the relationship between VO₂peak and imaging biomarkers varied by group, we conducted linear regression analyses in R. Each model included VO₂peak as the primary predictor, with age, sex, education, and group entered as covariates. A VO₂peak × group interaction term was added to assess whether the group moderated the association between VO₂peak and each biomarker. This allowed us to test whether the strength or direction of the VO₂peak–biomarker relationship differed between patients and controls. P-values were adjusted for multiple comparisons using the False Discovery Rate (FDR) procedure.

For regions in which the interaction term was statistically significant after FDR correction, we conducted post-hoc analyses by refitting the model separately within each group. This allowed us to estimate the VO₂_peak_–biomarker slope in patients and controls independently and identify which group was driving the observed interaction effect. To provide a complete picture of VO₂_peak_–biomarker relationships, we also performed these group-specific regressions across all regions regardless of interaction significance. However, these additional results are exploratory and are presented in the Supplementary Materials.

### Data availability

Anonymized and defaced data that support the findings of this study will be made openly available on OpenNeuro (https://openneuro.org) within six months of publication. A persistent DOI will be provided upon release.

## Results

### Cerebrovascular health and VO₂peak

In the full sample, VO₂peak was positively associated with both CBF and CVR, with no significant moderation by group. Higher VO₂peak was linked to increased CBF across the whole GM, including extensive regions of the frontal, parietal, and temporal cortices, as well as subcortical regions (Table 3; Figure 2). Similarly, CVR demonstrated a significant positive association with VO₂peak in global GM and in a more localized set of regions, primarily within the frontal, temporal, and precentral cortices. Detailed regional results are presented in Table 3 and Figure 2.

**Figure 2.**
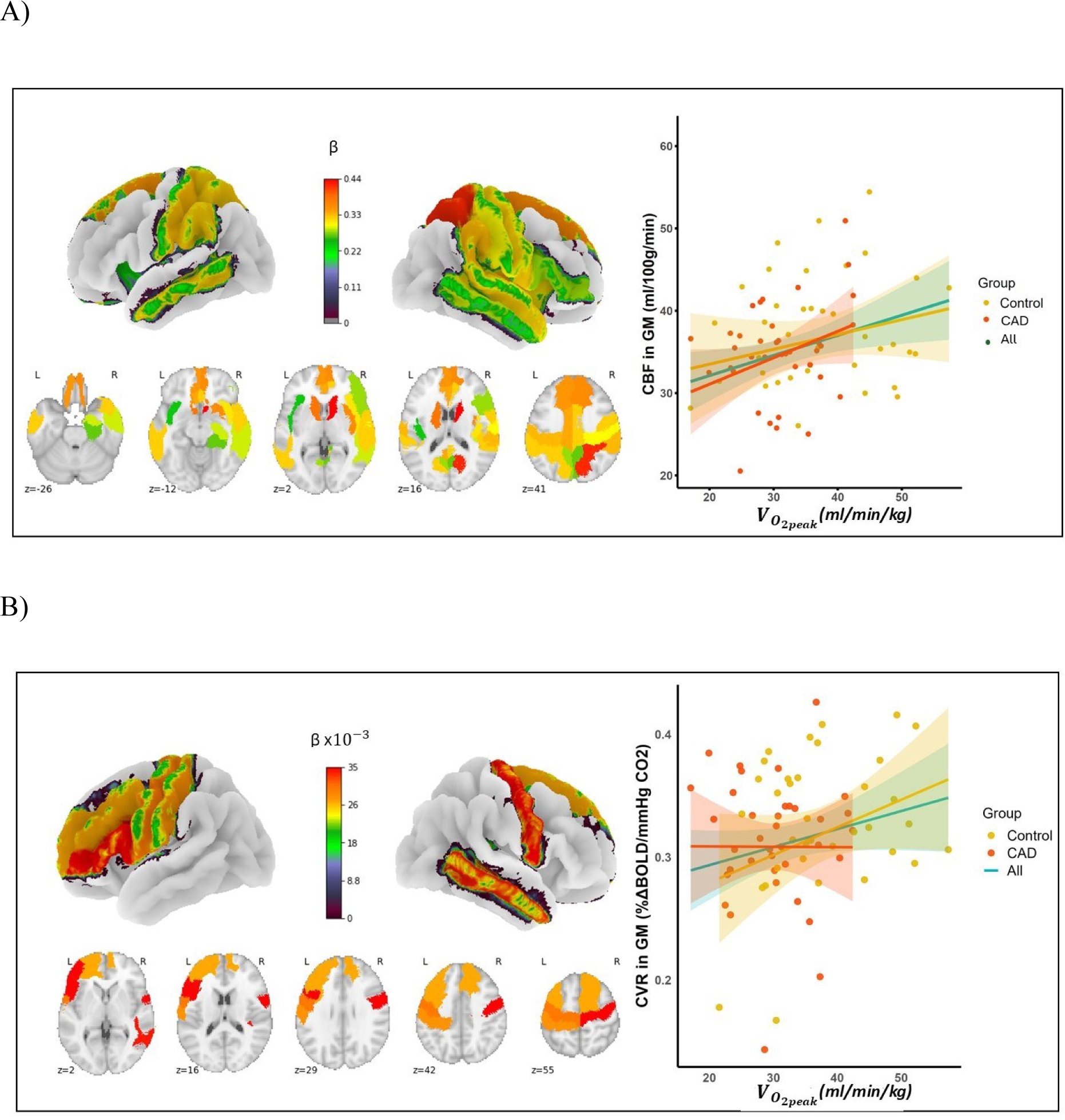
Associations of VO₂peak with CBF and CVR. Brain surface projections, on the left-hand side, display the beta values derived from linear regression analyses for all participants, adjusted for age, sex, and group as covariates. Only brain regions with significant associations are shown. The linear regression plot, on the right, illustrates the relationship between VO₂peak and CBF **(A)** and CVR **(B)** in the whole GM for all participants (green line) and each group separately (yellow for healthy controls and red for CAD).

**Table. 3.** Brain regions showing significant correlations between VO₂peak and CBF, CVR with age, sex and group as covariates.

|  | CBF |  |  | CVR |  |  |
| --- | --- | --- | --- | --- | --- | --- |
| | $\beta$ | P value | Adjusted-R<sup>2</sup> | $\beta$ | P value | Adjusted-R<sup>2</sup> |
| GM<sup>a</sup> | 0.32 | **0.02** | 0.17 | 0.002 | **0.04** | 0.10 |
| L Sup Frontal G | 0.35 | **0.03** | 0.15 | n. s | n. s | n. s |
| R Sup Frontal G | 0.36 | **0.03** | 0.13 | 0.002 | 0.04 | 0.15 |
| L Mid Frontal G | n. s | n. s | n. s | 0.003 | **0.04** | 0.16 |
| L Inf Frontal G | n. s | n. s | n. s | 0.003 | 0.04 | 0.10 |
| R Inf Frontal G | 0.28 | **0.02** | 0.12 | n. s | n. s | n. s |
| L Sup Parietal G | 0.32 | **0.03** | 0.13 | n. s | n. s | n. s |
| R Sup Parietal G | 0.42 | **<0.01** | 0.19 | n. s | n. s | n. s |
| L precuneus | 0.28 | **0.03** | 0.12 | n. s | n. s | n. s |
| R precuneus | 0.27 | **0.02** | 0.14 | n. s | n. s | n. s |
| R Sup Temporal G | 0.32 | **0.01** | 0.15 | n. s | n. s | n. s |
| L Sup Temporal G | n. s | n. s | n. s | n. s | n. s | n. s |
| R Mid Temporal G | 0.29 | **0.02** | 0.12 | 0.003 | **0.04** | 0.09 |
| L Mid Temporal G | 0.32 | **0.02** | 0.13 | n. s | n. s | n. s |
| R parahippocampus | 0.27 | **0.01** | 0.13 | n. s | n. s | n. s |
| R Insula | 0.31 | **<0.01** | 0.19 | n. s | n. s | n. s |
| L Insula | 0.24 | **0.03** | 0.15 | n. s | n. s | n. s |
| L Cingulate G | 0.34 | **0.01** | 0.15 | n. s | n. s | n. s |
| R cingulate G | 0.32 | **0.01** | 0.14 | n. s | n. s | n. s |
| L Caudate | 0.38 | **0.03** | 0.16 | n. s | n. s | n. s |
| R Caudate | 0.44 | **<0.01** | 0.18 | n. s | n. s | n. s |
| R Putamen | 0.36 | **<0.01** | 0.19 | n. s | n. s | n. s |
| R hippocampus | 0.29 | **0.01** | 0.13 | n. s | n. s | n. s |
| L precentral G | n. s | n. s | n. s | 0.003 | **0.04** | 0.12 |
| R precentral G | 0.32 | **0.03** | 0.14 | 0.003 | **0.04** | 0.13 |
| L postcentral G | 0.33 | **0.03** | 0.13 | 0.002 | **0.04** | 0.10 |
| R postcentral G | 0.30 | **0.03** | 0.15 | n. s | n. s | n. s |
| L Sup Marginal G | 0.32 | **0.02** | 0.12 | n. s | n. s | n. s |
| R Sup Marginal G | 0.33 | **0.01** | 0.15 | n. s | n. s | n. s |

<sup>a</sup> GM: Grey Matter; L: left ; R: right; Sup: superior; Mid: Middle; G: gyrus.

### Cerebral metabolic health and VO₂peak

Although no significant association between VO₂peak and either OEF or CMRO₂ was observed when analyzing the overall sample, moderation analysis revealed that group significantly moderated both relationships. Specifically, the VO₂peak × group interaction was significant for multiple brain regions in both OEF and CMRO₂ models, indicating that the association between cardiorespiratory fitness and these brain biomarkers differed between patients and healthy controls.

To further explore the nature of these interactions, post-hoc linear models were conducted separately within each group. For CMRO₂, a positive association with VO₂peak was observed only in the CAD group, with effects in whole GM and several grey matter regions including the left middle temporal gyrus, left superior temporal gyrus, left inferior temporal gyrus, and the left and right middle orbital gyri, as well as the middle frontal gyri. In contrast, for OEF, a negative association with VO₂peak was observed only in the healthy control group. Significant effects were found in the GM, left middle temporal gyrus, as well as in the superior and inferior temporal gyri, middle orbital gyrus, and middle frontal gyri. Full statistical results for moderation analysis are reported in Table 4 and can be visualized in Figure 3. Post-hoc analyses results are reported in Table 5.

**Figure 3.**
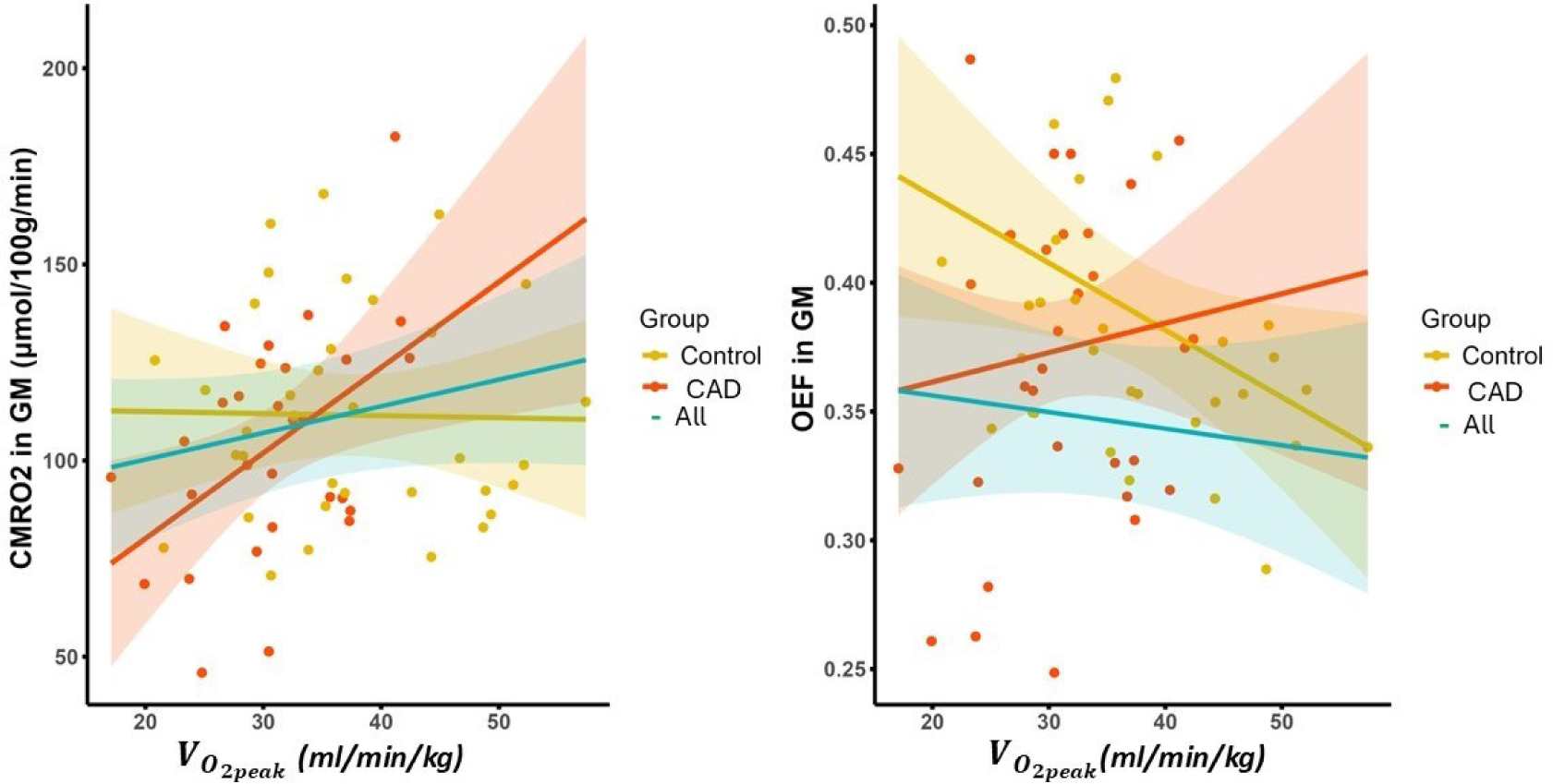
Associations between VO₂peak (of lean body mass) and brain metabolic markers in GM, in all participants and in each group. **(A)** Relationship between VO₂peak and CMRO₂ showing a significant positive association in the CAD group (red line) but not in controls (yellow line) or when analyzing all participants together (green line). **(B)** Relationship between VO₂peak and OEF showing a negative association in the control group (yellow line) compared to a null relationship in CAD (red line) and the overall group (green dashed line). Shaded areas represent 95% confidence intervals. These results illustrate distinct group-dependent patterns linking cardiorespiratory fitness to brain metabolism.

**Table. 4.**
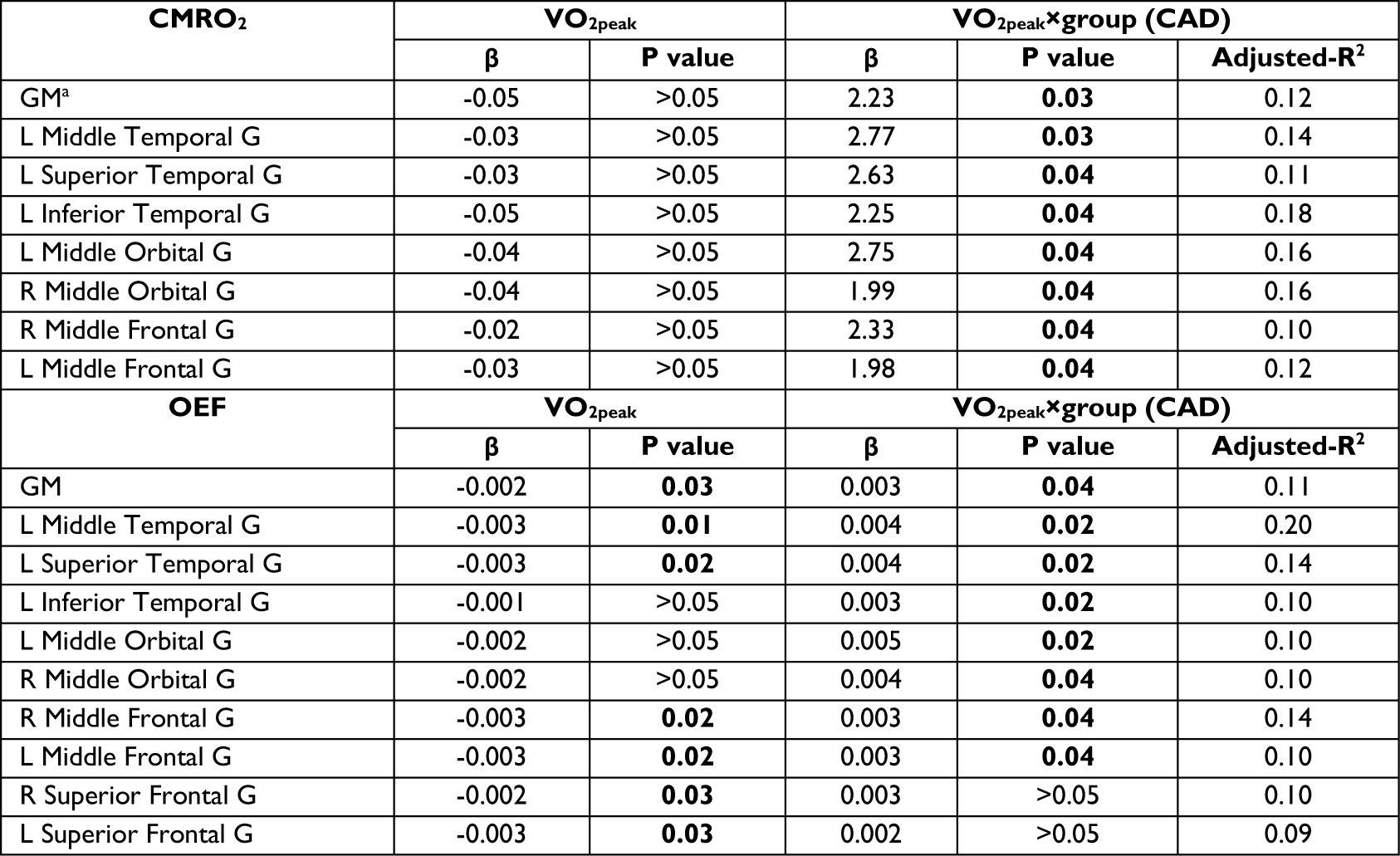
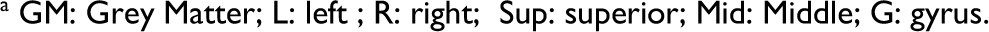
Regions showing significant group moderation of VO₂peak–OEF and VO₂peak–CMRO₂ associations (CAD vs. controls)

| CMRO<sub>2</sub> | VO<sub>2</sub>peak | | VO<sub>2</sub>peak×group (CAD) | | |
| --- | --- | --- | --- | --- | --- |
| | $\beta$ | P value | $\beta$ | P value | Adjusted-R<sup>2</sup> |
| GM<sup>a</sup> | -0.05 | >0.05 | 2.23 | **0.03** | 0.12 |
| L Middle Temporal G | -0.03 | >0.05 | 2.77 | **0.03** | 0.14 |
| L Superior Temporal G | -0.03 | >0.05 | 2.63 | **0.04** | 0.11 |
| L Inferior Temporal G | -0.05 | >0.05 | 2.25 | **0.04** | 0.18 |
| L Middle Orbital G | -0.04 | >0.05 | 2.75 | **0.04** | 0.16 |
| R Middle Orbital G | -0.04 | >0.05 | 1.99 | **0.04** | 0.16 |
| R Middle Frontal G | -0.02 | >0.05 | 2.33 | **0.04** | 0.10 |
| L Middle Frontal G | -0.03 | >0.05 | 1.98 | **0.04** | 0.12 |
| OEF | VO<sub>2</sub>peak | | VO<sub>2</sub>peak×group (CAD) | | |
| | $\beta$ | P value | $\beta$ | P value | Adjusted-R<sup>2</sup> |
| GM | -0.002 | **0.03** | 0.003 | **0.04** | 0.11 |
| L Middle Temporal G | -0.003 | **0.01** | 0.004 | **0.02** | 0.20 |
| L Superior Temporal G | -0.003 | **0.02** | 0.004 | **0.02** | 0.14 |
| L Inferior Temporal G | -0.001 | >0.05 | 0.003 | **0.02** | 0.10 |
| L Middle Orbital G | -0.002 | >0.05 | 0.005 | **0.02** | 0.10 |
| R Middle Orbital G | -0.002 | >0.05 | 0.004 | **0.04** | 0.10 |
| R Middle Frontal G | -0.003 | **0.02** | 0.003 | **0.04** | 0.14 |
| L Middle Frontal G | -0.003 | **0.02** | 0.003 | **0.04** | 0.10 |
| R Superior Frontal G | -0.002 | **0.03** | 0.003 | >0.05 | 0.10 |
| L Superior Frontal G | -0.003 | **0.03** | 0.002 | >0.05 | 0.09 |

<sup>a</sup> GM: Grey Matter; L: left ; R: right; Sup: superior; Mid: Middle; G: gyrus.

**Table 5.** Post-hoc linear regression results of the association between VO₂peak and brain metabolic biomarkers (CMRO₂ and OEF) within each group separately.

| CMRO<sub>2</sub> in CAD | VO<sub>2peak</sub> |  |  |
| --- | --- | --- | --- |
| | $\beta$ | P value | Adjusted-R<sup>2</sup> |
| GM<sup>a</sup> | 2.37 | 0.03 | 0.22 |
| L Middle Temporal G | 2.47 | 0.03 | 0.21 |
| L Superior Temporal G | 2.40 | 0.04 | 0.16 |
| L Inferior Temporal G | 2.16 | 0.03 | 0.23 |
| L Middle Orbital G | 2.35 | 0.04 | 0.14 |
| R Middle Orbital G | 2.14 | 0.04 | 0.13 |
| R Middle Frontal G | 1.87 | 0.03 | 0.19 |
| L Middle Frontal G | 1.75 | 0.04 | 0.12 |
| OEF in HC | VO<sub>2peak</sub> |  |  |
| | $\beta$ | P value | Adjusted-R<sup>2</sup> |
| GM | -0.003 | <0.01 | 0.29 |
| L Middle Temporal G | -0.003 | <0.01 | 0.28 |
| L Superior Temporal G | -0.003 | <0.01 | 0.25 |
| L Inferior Temporal G | -0.001 | 0.04 | 0.09 |
| L Middle Orbital G | -0.003 | 0.03 | 0.15 |
| R Middle Orbital G | -0.003 | <0.01 | 0.16 |
| R Middle Frontal G | -0.004 | 0.03 | 0.27 |
| L Middle Frontal G | -0.003 | 0.03 | 0.15 |

<sup>a</sup> GM: Grey Matter; L: left ; R: right; Sup: superior; Mid: Middle; G: gyrus.

## Discussion

In this study, we examined the associations between VO₂_peak_, a measure of cardiorespiratory fitness, and critical cerebral vascular and metabolic biomarkers in healthy controls and individuals with CAD. Our analysis revealed significant positive relationships between VO₂_peak_ and both CBF and CVR, across the entire study cohort. These findings emphasize the beneficial effects of higher cardiorespiratory fitness on cerebrovascular health across the cardiovascular health spectrum. In contrast, no significant associations were observed between VO₂_peak_ and either CMRO₂ or OEF in analyses combining both groups. However, moderation analyses revealed distinct associations in CAD patients compared to controls. We found VO₂_peak_ to be associated with greater CMRO₂ in CAD participants within gray matter regions of the temporal lobes and orbital frontal cortex. This suggests that enhanced fitness may be beneficial for metabolic oxygen utilization in these brain regions, especially in individuals with CAD. Conversely, fitness was found to be associated with lower OEF exclusively in healthy controls within temporal and frontal gray matter regions. This result implies that in healthy individuals, the effects of cardiorespiratory fitness are predominantly reflected in higher CBF, with an unchanged CMRO_2_, leading to lower OEF. This results in greater oxygen availability at rest. In contrast, in patients, concurrent improvements in CMRO_2_ and CBF cancel out to result in unchanged OEF.

We observed widespread positive associations between VO₂_peak_ and CBF in frontal, parietal, temporal, and key subcortical regions (insula, hippocampus, caudate, putamen). This pattern echoes previous findings in CAD cohorts (MacIntosh et al., 2014) and in several studies in healthy participants showing higher CBF in grey matter and the hippocampus with greater fitness (Chaddock-Heyman et al., 2016; Upadhyay et al., 2022; Zimmerman et al., 2014). It is interesting to note however, that these findings may reflect the physiological processes at the normal or lower range of fitness, since some studies in very highly trained adults report null or negative relationships (Intzandt et al., 2020; Krishnamurthy et al., 2022; Olivo et al., 2021; Van Praag et al., 2005). These discrepancies in highly fit adults are potentially due to a ceiling effect or greater oxygen extraction efficiency, which may reduce the need for high resting perfusion (Furby et al., 2020; Olivo et al., 2021; Van Praag et al., 2005). In our sample, the positive association may reflect fitness-related vascular adaptations, including increased capillary density shown in animal studies (Kleim et al., 2002; Swain et al., 2003) and enhanced perfusion in metabolically active regions.

Our CVR findings add important nuance to the existing literature by demonstrating a positive relationship between VO₂_peak_ normalized by lean body mass (LBM), and CVR across both healthy individuals and patients with CAD. Specifically, significant associations were identified in key cortical regions such as the superior and middle frontal gyri, parietal lobes, middle temporal gyrus, precentral gyrus, and supramarginal gyrus—areas particularly susceptible to aging-related changes and known targets for exercise-induced neuroplasticity (Chen et al., 2011; d’Arbeloff, 2020). This positive relationship aligns with prior findings showing aerobic exercise’s beneficial impacts on cerebrovascular health, likely reflecting vascular adaptations such as increased arterial compliance and enhanced endothelial function (Tan et al., 2017; Tanaka et al., 2001; Vaitkevicius et al., 1993). Regular aerobic activity is known to induce structural remodeling and biochemical improvements, including upregulated endothelial nitric oxide production and reduced oxidative stress (Barnes et al., 2021; Thijssen et al., 2016), likely collectively enhancing vasodilatory capacity and CVR.

Nevertheless, our results should be interpreted within the context of varied literature findings. Many of the previously documented positive relationships between CVR and VO₂_peak_ were obtained by measuring CVR with trans-cranial Doppler imaging (Barnes et al., 2013; Bliss et al., 2023; Braz et al., 2017; Murrell et al., 2013). MRI-based studies investigating the relationship between cardiorespiratory fitness and CVR have produced inconsistent findings, and our study is among the first to demonstrate a positive association. Previous studies using MRI have reported negative (Intzandt et al., 2020; Penukonda et al., 2025; Potvin-Jutras et al., 2024), quadratic (DuBose et al., 2022), and null relationship (Flück et al., 2014). Most notably, Intzandt and colleagues (2020), whose methodology closely resembles ours (using hypercapnic BOLD-CVR and VO₂_peak_), reported negative associations between fitness and both CVR and CBF in healthy older adults. The critical difference between their study and ours is the study population; whereas Intzandt included only healthy older individuals, our study comprised a mixed group of CAD patients and controls. The positive association we observed may therefore reflect a particularly beneficial impact of cardiorespiratory fitness on cerebrovascular function within CAD patients—a notion consistent with previous evidence showing improved cerebral perfusion following cardiac rehabilitation in similar populations (Anazodo et al., 2016). Additionally, differences in participant demographics, particularly sex composition, could contribute to this discrepancy. Our sample included a smaller proportion of females (22%), while Intzandt et al. included 66% females, which is important given prior evidence of sex-specific cerebrovascular responses to physical activity (Potvin-Jutras et al., 2024; Sabra et al., 2021). Other studies such as (DuBose et al., 2022) further emphasize the complexity of the CVR–fitness relationship. DuBose et al. proposed an inverted-U (quadratic) relationship, suggesting optimal CVR at moderate fitness levels. Collectively, these varied outcomes highlight the nuanced, population-dependent nature of the fitness–CVR association, emphasizing the need to consider population characteristics and methodological factors when interpreting findings across studies.

Notably, although no significant relationship between VO₂_peak_ and CMRO₂ was found in the overall study population, we observed a group interaction, showing that group moderates the relationship between fitness and oxygen usage. More specifically, VO₂_peak_ was positively related to CMRO₂ within the CAD group across gray matter, particularly in frontal and temporal regions—areas known to be susceptible to the effects of aging and vascular disease (Chen et al., 2011; Ovsenik et al., 2021). This association represents a novel finding exclusive to patients with CAD. Our results contrast with a recent study (Huppert and Qiao, 2025) which reported a negative relationship between global CMRO₂ and VO₂_peak_ in healthy older adults. This discrepancy could be due to differences in participant demographics, especially given that the same group of individuals with CAD were previously shown to have CMRO_2_ deficits (Sanami et al., 2025). Alternatively, some of these discrepancies could be due to different methodological approaches. Gay et al. measured global CMRO₂ using TRUST, a T2-based method. Studies using TRUST have typically shown increased CMRO_2_ with aging (Lu et al., 2011; Peng et al., 2014), in contrast to the decrease documented with the T2*-based approaches such as the one used here (De Vis et al., 2015). Because all other parameters tested are improved with fitness in our healthy sample, our findings suggest that CMRO₂ may be at a ceiling in healthy older adults and only when CMRO₂ is affected by disease can it be improved through higher cardiorespiratory fitness. In CAD, the positive effect of VO₂_peak_ on CMRO₂ might indicate improved mitochondrial function, which plays a central role in oxygen metabolism and is known to be compromised at least outside the brain in CAD (Mengozzi et al., 2025).

To our knowledge, this is the first study to investigate the effect of VO₂_peak_ on OEF and CMRO_2_, in a population that includes both healthy individuals and patients with CAD. The negative association between OEF and VO₂_peak_, observed in healthy participants across multiple brain regions including the whole grey matter, suggests a compensatory mechanism in response to lower cardiorespiratory fitness. In healthy individuals with lower VO₂_peak_, the brain compensates for reduced oxygen delivery by increasing OEF, extracting more oxygen from the blood to meet its metabolic demands and maintain CMRO_2_. This indicates that cerebral oxygen metabolism remains stable despite reduced fitness levels due to the brain’s ability to adjust oxygen extraction. In contrast, the absence of a significant relationship between OEF and VO₂_peak_ in patients implies that this compensatory mechanism may be exhausted, having already led to metabolic impairment either through cell loss or mitochondrial dysfunction. This is consistent with our previous results (Sanami et al., 2025) showing lower CMRO₂ in individuals with CAD.

The negative relationship between OEF and VO₂_peak_ in healthy individuals suggests enhanced oxygen extraction to compensate for lower cardiorespiratory fitness. The absence of this relationship in CAD patients indicates compromised compensatory capacity, highlighting the importance of fitness interventions to maintain cerebral metabolic integrity. These findings suggest a continuum of adaptive responses, where metabolic flexibility diminishes with disease progression, but can be rescued by higher cardiorespiratory fitness.

## Limitations

The imaging technique used in this study, arterial spin labeling (ASL), is inherently limited by low temporal signal-to-noise ratio (tSNR), which can be further reduced in populations with vascular risk due to prolonged arterial transit times (the time taken for labeled blood to travel from labeling to imaging location). The use of a fixed post-labeling delay (PLD)—the waiting time between blood labeling and image acquisition—may have led to an underestimation of CBF in individuals with delayed blood transit. Future studies should consider acquiring data at multiple PLDs to enhance the accuracy of CBF measurements. Additionally, background suppression techniques, which were not applied in this dual-echo acquisition to preserve the BOLD signal for estimation of CVR, OEF, and CMRO₂, could improve ASL signal quality. Further limitations include the homogeneity of our sample, recruited from a single site, predominantly male, and excluding individuals with severe CAD. This significantly limits the generalizability of our findings. Moreover, our sample was also highly homogeneous in terms of socioeconomic status, education, and ethnicity. Future studies should thus recruit broader and more diverse cohorts to validate and expand upon our findings.

Additionally, the absence of direct measures of cardiac function restricts interpretation of the mechanisms linking cardiovascular performance to cerebrovascular health. Integrating cardiac functional assessments in future studies would better clarify heart–brain interactions. Finally, this study utilized cross-sectional measurements, which precludes causal inference. Intervention studies—particularly those exploring whether exercise interventions in CAD populations can improve these cerebrovascular and metabolic markers—represent an essential next step to investigate underlying mechanisms more robustly.

## Conclusion

This study provides novel evidence linking cardiorespiratory fitness, measured as VO₂_peak_, with both vascular and metabolic markers of brain health in older adults with and without coronary artery disease. We demonstrate widespread positive associations between VO₂peak and CBF and CVR, suggesting that aerobic fitness supports vascular function regardless of the presence of cardiovascular pathology. Importantly, we also show that metabolic responses to aerobic fitness differ between healthy individuals and CAD patients. Patients showed a positive association between VO₂peak and CMRO₂, particularly in brain regions vulnerable to aging and disease, indicating that higher aerobic fitness can help preserve oxidative metabolism, either through improved mitochondrial function, neurogenesis, or other neurotrophic mechanisms. In contrast, healthy participants exhibited an elevated OEF with lower fitness, likely as a compensatory mechanism to preserve CMRO₂ despite lower CBF. These findings underscore the potential of VO₂peak as a modifiable factor influencing both vascular and metabolic resilience in the brain, both in patients with heart disease and in older individuals without known vascular risk factors or disease. Moreover, they suggest that targeted aerobic interventions may hold value not only for systemic cardiovascular health, but also for preserving or enhancing brain function, especially in populations with cardiac disease who are at elevated risk of cognitive decline and stroke.

## Acknowledgements

We would like to thank everyone who contributed to this project: Paule Samson, Thomas Vincent, Julie Lalongé, Hakima Benhalima, Milla Shakleva, Victoria D’Amours, Agathe Godet, Stephanie Beram, Roni Zaks, Robert Hovey, Alexandre Bailey, Catherina Medeiros, Amélie Mainville-Berthiaume and Zineb Rouabah. Thank you also to the laboratories of Dr Louis Bherer and Dr Mathieu Gayda. Lastly, we would like to acknowledge our research participants without whom none of this would have been possible.

## Funding

This work was supported by funding awarded to Claudine J. Gauthier from the Natural Sciences and Engineering Research Council of Canada (NSERC Discovery Grant: RGPIN-2015-04665; 2024-06455), Fonds de recherche du Québec (FRQ 5232), the Heart and Stroke Foundation of Canada (G-17-0018336), the Heart and Stroke Foundation New Investigator Award, the Henry J.M. Barnett Scholarship, the Michal and Renata Hornstein Chair in Cardiovascular Imaging and the Mirella and Lino Saputo research chair in cardiovascular health and the prevention of cognitive decline. Additional support was provided by the Canadian Institutes of Health Research (FRN: 175862, to Stefanie A. Tremblay), the Vascular Training Platform (VAST) (to Ali Rezaei), the Heart and Stroke Foundation of Canada and Brain Canada (to Zacharie Potvin-Jutras), and the Alzheimer Society Research Award (to Brittany Intzandt.).

## Competing interests

The authors report no competing interests.

## Notes

### Competing Interest Statement

The authors have declared no competing interest.

